# Presaccadic attention enhances contrast sensitivity, but not at the upper vertical meridian

**DOI:** 10.1101/2021.10.01.461760

**Authors:** Nina M. Hanning, Marc M. Himmelberg, Marisa Carrasco

## Abstract

Human visual performance is not only better at the fovea and decreases with eccentricity, but also has striking radial asymmetries around the visual field: At a fixed eccentricity, it is better along (1) the horizontal than vertical meridian and (2) the lower than upper vertical meridian. These asymmetries are not alleviated by covert exogenous or endogenous attention, but have been studied exclusively during eye fixation. However, a major driver of everyday attentional orienting is saccade preparation, during which visual attention automatically shifts to the future eye fixation. This presaccadic shift of attention is considered strong and compulsory, and relies on fundamentally different neural computations and substrates than covert attention. Given these differences, we investigated whether presaccadic attention can compensate for the ubiquitous performance asymmetries observed during eye fixation. Our data replicate polar performance asymmetries during fixation and document the same asymmetries during saccade preparation. Crucially, however, presaccadic attention enhanced contrast sensitivity at the horizontal and lower vertical meridian, but not at the upper vertical meridian. Thus, instead of attenuating polar performance asymmetries, presaccadic attention exacerbates them.

## Introduction

Human visual performance is asymmetric around the visual field. At isoeccentric locations it is better along the horizontal than vertical meridian (horizontal-vertical anisotropy; HVA) and along the lower than upper vertical meridian (vertical-meridian asymmetry; VMA) (e.g., Mackeben 1999; Altpeter et al., 2000; Carrasco et al., 2001; Cameron et al., 2002; Baldwin et al., 2012; Greenwood et al., 2017; Himmelberg et al., 2020; Barbot et al., 2021). These polar angle asymmetries, known as *performance fields*, are pervasive –they emerge for many visual dimensions (e.g., contrast sensitivity, spatial resolution), regardless of head rotation, stimulus orientation or display luminance, and whether measured monocularly of binocularly (e.g., Carrasco et al. 2001; Corbett & Carrasco, 2012; Barbot et al., 2021). Moreover, performance fields are resilient, they are not alleviated by spatial covert attention, deployed in the absence of eye movements: Both *exogenous* attention, which is involuntary, fast and transient (Carrasco et al., 2001; Cameron et al., 2002; Roberts et al., 2016, 2018; Talgar & Carrasco, 2002) and *endogenous* attention, which is voluntary, slower and sustained (Purokayastha et al., 2021), uniformly improve performance around the visual field, thereby preserving the polar angle asymmetries (***Figure 1A***).

**Figure 1.**
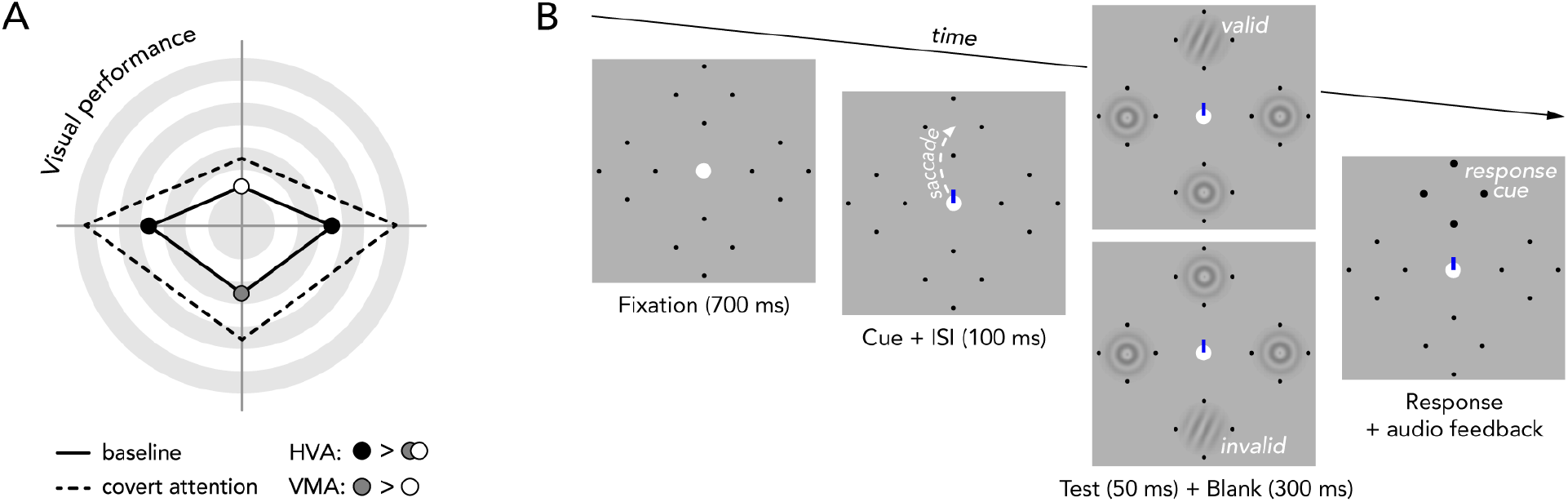
Background and experimental design. (**A**) Schematic representation of polar angle performance asymmetries reported in previous research. Points further from the center represent higher performance. At matched eccentricity, visual performance is better on the horizontal than vertical meridian (HVA), and along the lower than upper vertical meridian (VMA) in several fundamental perceptual dimensions, including contrast sensitivity. (**B**) Experimental task. After a fixation period, a central direction cue (blue line) appeared. Observers were instructed to make a saccadic eye movement to the indicated target (8.5° left, right, above, or below central fixation, marked by black dots). Note that the dashed arrow indicates saccade preparation. 100 ms after direction cue onset, a test Gabor patch was presented at either the saccade target (*valid*) or at the opposing isoeccentric location (*invalid*). Radial frequency patterns with no orientation information were presented at the other three locations. At the end of the trial (after saccade offset), a response cue (bolded placeholder) indicated the location at which the test Gabor had appeared, and observers reported the Gabor orientation. Note that in the fixation condition the cue pointed to two opposing potential test locations (left and right or upper and lower) (supplemental ***Video S1*** demonstrates the trials sequence of each condition).

Presaccadic attention, which automatically shifts to the future eye fixation during the preparation of saccadic eye movements also benefits visual perception: Already before saccade onset, performance at the saccade target is enhanced, which is, for example, reflected in improved letter and orientation discrimination accuracy (Kowler et al., 1995; Deubel & Schneider, 1996; Montagnini & Castet, 2007). However, covert and presaccadic attention differentially modulate visual perception and the representation of features of basic visual dimensions, such as orientation and spatial frequency (Collins et al., 2010; Li et al., 2016; Ohl et al., 2017). They engage different neural computations –contrast gain and response gain (Li et al., 2021b)– and recruit partially distinct neural populations in frontal eye fields (FEF) and superior colliculus (SC) that control covert attention and saccade preparation in human and nonhuman primates (e.g., Ignashchenkova et al., 2004; Gregoriou et al., 2012; Messinger et al., 2021).

Here we investigated whether presaccadic attention, unlike covert exogenous and endogenous attention, can compensate for the polar angle performance asymmetries established during eye fixation. Presaccadic attention is considered to be deployed automatically, yet there are some implicit cognitive processes that can permeate its effects and help distribute resources to potential relevant locations (White et al., 2013). Given the prevalence of presaccadic attention in the selective processing of visual information and its differences with covert attention (review: Li et al., 2021b), we hypothesized that presaccadic attention may benefit performance more where it is worse (i.e., at the upper vertical meridian), and thus diminish polar performance asymmetries. For this reason, we separately assessed presaccadic benefits (at the saccade target) and costs (at the opposite location) along the horizontal and vertical meridians, compared to a fixation baseline condition.

Our data reproduced both the HVA and VMA during fixation (e.g., Carrasco et al., 2001; Cameron et al., 2002; Abrams et al., 2012; Roberts et al., 2018; Himmelberg et al., 2020; Purokayastha et al., 2021) and reveal the same polar asymmetries for contrast sensitivity during saccade preparation. Crucially, contrary to our initial hypothesis, presaccadic attention did not attenuate the cardinal polar angle asymmetries: It enhanced contrast sensitivity at the horizontal and lower vertical meridian, but *not* at the upper vertical meridian. The surprising absence of a performance advantage preceding upwards saccades suggests a rigid perceptual limitation along the upper vertical meridian that cannot be allayed by presaccadic attention.

## Results

We measured contrast sensitivity during saccade preparation using a two-alternative forced-choice orientation discrimination task (***Figure 1B***). Eleven observers performed horizontal or vertical saccades to a centrally cued peripheral target (8.5°eccentricity) and discriminated the orientation of a ±15° tilted Gabor grating presented briefly, just before eye movement onset, either at the saccade target (*valid*) or the opposite (*invalid*) isoeccentric location. In a fixation condition (*baseline*), observers performed the same orientation discrimination task without preparing saccades.

For each experimental condition (*valid*, *invalid*, *baseline*) we independently titrated Gabor contrast at four polar angle locations (*left, right, upper, lower*) using an adaptive psychometric “staircase” procedure (see Materials and methods - Titration procedure) and computed contrast sensitivity as the reciprocal of the titrated threshold value (***Figure 2A***). Below we report Greenhouse-Geisser corrected p-values for ANOVAs for which the assumption of sphericity was not met. All post-hoc comparisons are Bonferroni corrected for multiple comparisons.

**Figure 2.**
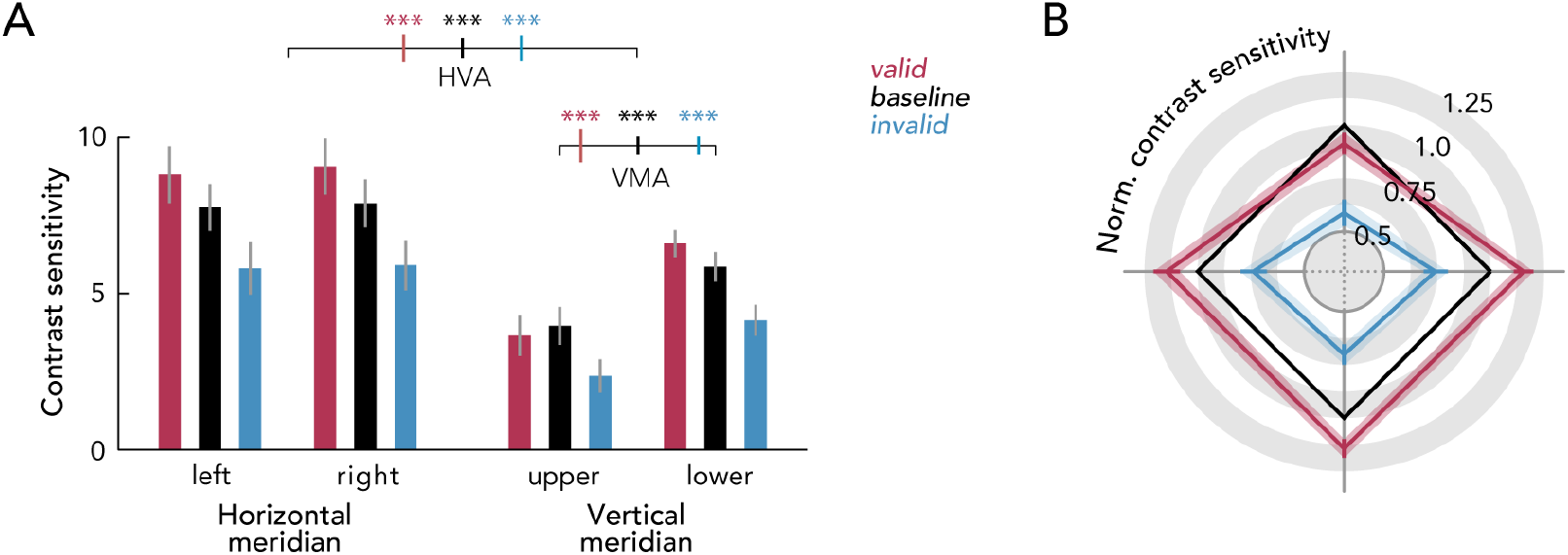
Main results. (**A**) Contrast sensitivity as a function of cardinal location for each experimental condition. Error bars depict ±1 standard error of the mean (SEM). Horizontal brackets indicate horizontal vs. vertical meridian (HVA) and vertical meridian upper vs. lower (VMA) comparisons, color coded error bars on the brackets indicate ±1 standard error of the difference between the compared conditions (SED). \**p*<0.05, \*\**p*<0.01, \*\*\**p*<0.001. (**B**) Relative contrast sensitivity (normalized by respective baseline sensitivity) for each condition and cardinal location. Shaded error areas indicate ±1 SEM.

To test for the HVA, we conducted a 3 (condition: valid, invalid, baseline) X 2 (meridian: horizontal, vertical) repeated-measures ANOVA. The main effects for condition (*F*(2,20)=41.78, *p*<0.001) and meridian (*F*(1,10)=74.51, *p*<0.001), and their interaction (*F*(2,20)=8.45, *p*=0.004), were significant. Likewise, to test for the VMA, a 3 (condition: valid, invalid, fixation) X 2 (vertical meridian: upper, lower) repeated-measures ANOVA yielded significant main effects for condition (*F*(2,20)=44.18, *p*<0.001) and location (*F*(1,10)=42.46, *p*<0.001), and a significant interaction (*F*(2,20)=7.46, *p*=0.018). Post-hoc comparisons confirmed that for the fixation condition, contrast sensitivity changed around the visual field yielding the HVA (horizontal-vertical difference: 2.91±0.35 mean±SEM, *p*<0.001) and VMA (upper-lower vertical difference: 1.89±0.30, *p*<0.001), consistent with previous findings (Carrasco et al., 2001; Cameron et al., 2002; Abrams et al., 2012; Himmelberg et al., 2020; Purokayastha et al., 2021). Importantly, the same asymmetries emerged during saccade preparation, both when tested at the saccade target (*valid*) and opposite of it (*invalid*): HVA (*valid*: 3.81±0.47, *p*<0.001, *invalid*: 2.61±0.44, *p*<0.001) and VMA (*valid*: 2.95±0.56, *p*<0.001, *invalid*: 1.78±0.30, *p*<0.001).

To visualize the interaction between experimental condition and location, we plotted contrast sensitivity as a ratio of the baseline condition (***Figure 2B***). The 2 (condition: valid, fixation) X 4 (location: left, right, upper, lower) repeated-measures ANOVA to evaluate the relative benefit of presaccadic attention over the baseline yielded significant main effects for condition (*F*(1,10)=7.05, *p*=0.024), location (*F*(3,30)=7.24, *p*=0.001), and their interaction (*F*(3,30)=7.24, *p*=0.001). Remarkably, post-hoc comparisons revealed the well-established perceptual advantage caused by presaccadic attention for all but one location: the upper vertical meridian. Relative to fixation (*baseline*), the preparation of horizontal and downward saccades increased contrast sensitivity at the target of the upcoming eye movement (*left*: *p*=0.040; *right*: *p*=0.007; *downward*: *p*=0.008). However, upward saccades did not yield a sensitivity benefit (*p*=0.114), and this effect was consistent across individual observers (***Figure 3A***).

**Figure 3.**
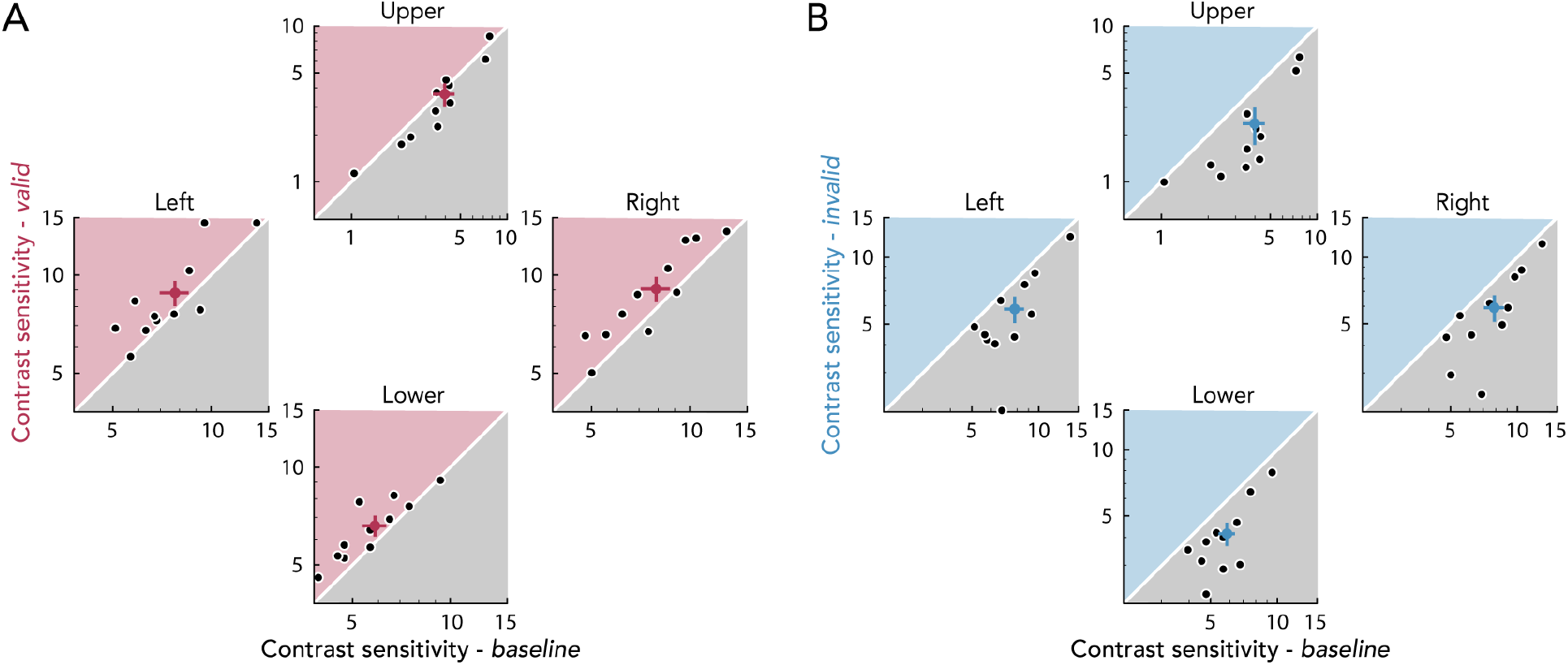
Individual observers’ presaccadic benefits and costs for each cardinal location. (**A**) Benefits. Individual observers’ sensitivity in valid trials plotted against their baseline sensitivity. (**B**) Costs. Individual observers’ sensitivity in invalid trials plotted against their baseline sensitivity. White dots with black lines depict the group average and ±1 SEM. Note that the axes are in log-scale and that the range for the upper panels differs to account for lower contrast sensitivity at the upper vertical meridian.

All locations, however, showed a pronounced cost when a saccade was prepared away from the test Gabor (***Figure 3B***). The 2 (condition: invalid, fixation) X 4 (location: left, right, upper, lower) repeated-measures ANOVA to evaluate the relative cost in performance below the baseline showed significant main effects for condition (*F*(1,10)=43.40, *p*<0.001), location (*F*(3,30)=3.64, *p*=0.038), and their interaction (*F*(3,30)=3.64, *p*=0.038). Post-hoc comparisons revealed a decrease in contrast sensitivity at the location opposite to the saccade target for all four tested locations (*left*: *p*=0.002; *right*: *p*<0.001; *downward*: *p*<0.001; and *upward*: *p*<0.001). To explore whether the local effect of presaccadic attention was related to the magnitude of participants’ visual field asymmetries during fixation, we separately assessed the correlations between the benefits and costs at each of the tested locations and the strength of HVA and VMA (see supplemental ***Table S2***), but none of the correlations were significant.

We evaluated eye movement parameters (***Figure 4***) to rule out that the absence of a performance advantage preceding upwards saccades can be explained by differences in saccade precision or latency. For example, the missing presaccadic benefit at the upper vertical meridian may be caused by comparatively slower or less precise upward saccade execution. A repeated-measures ANOVA found a significant main effect of saccade direction (*F*(3,30)=10.06, *p*<0.001) on latency (***Figure 4A***). Post-hoc comparisons showed that saccadic reaction times were significantly slower for downward than upward saccades (*p*=0.007). This effect is consistent with previous research in human (Honda & Findley, 1992; Tzelepi et al., 2010) and nonhuman (Schlykowa et al., 1996; Zhou & King, 2002) primates. A repeated-measures ANOVA showed a significant main effect of saccade direction (*F*(3,30)=6.99, *p*=0.005) on amplitude. Post-hoc comparisons indicated that saccade amplitudes (***Figure 4B***) were significantly shorter for upward than leftward (*p*=0.041) and rightward (*p*=0.041) saccades. However, consistent with previous work (Deubel & Schneider, 1996; Van der Stigchel & de Vries, 2015; Wollenberg et al., 2018; Hanning et al., 2019b), the presaccadic benefit was unaffected by landing precision (***Figure 4C***). For the upper vertical meridian, more precise saccades (landing closer to the target center) were not more likely to contribute to a staircase contrast decrement than less precise saccades (*p*=0.837; ***Figure 5A***, upper plot). Likewise, staircase contrast was not more likely to decrease with faster than slower saccade latency trials (*p*=0.346; ***Figure 5B***, upper plot). The missing presaccadic benefit at the upper vertical meridian, therefore, cannot be explained by differences in eye movement parameters among polar angle locations. Had saccade precision or latency affected the presaccadic attention benefit, trials with smaller/larger landing errors or faster/slower saccade latencies would have differentially affected staircase direction. Note that the same latency and precision comparison also did not yield significant differences for the remaining three tested locations (***Figure 5A-B** left, right*, and *lower* plots; all *p*>0.05).

**Figure 4.**
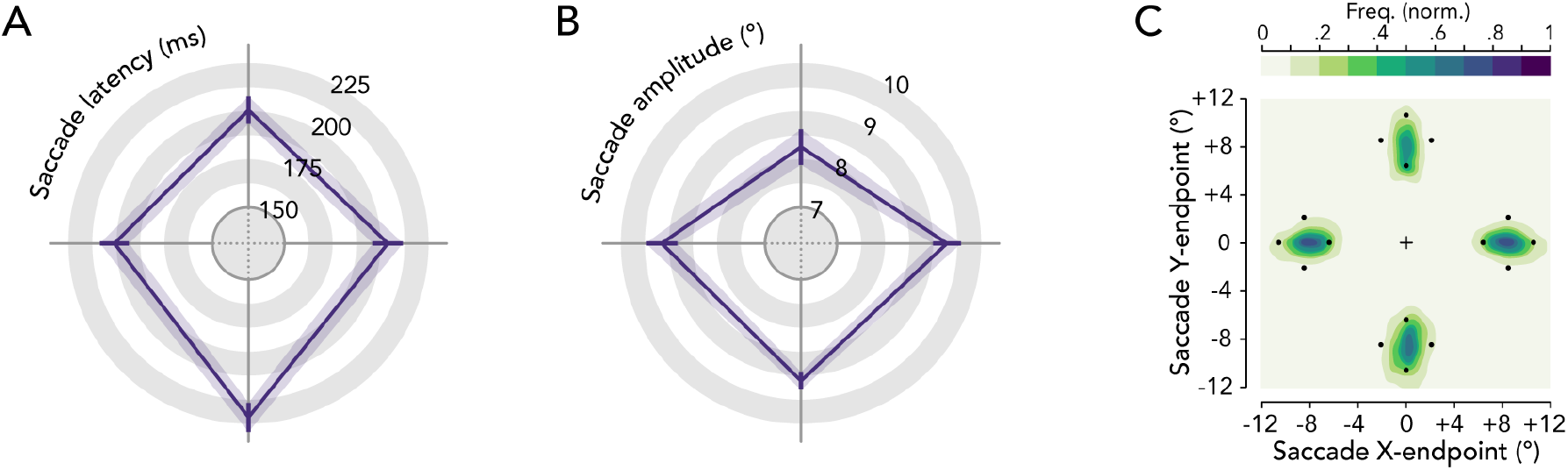
Eye movement parameters. Group average saccade latency (**A**) and saccade amplitude (**B**) as a function of saccade direction. Shaded error areas indicate ±1 SEM. (**C**) Normalized saccade endpoint frequency maps averaged across participants depicting saccade landing variance.

**Figure 5.**
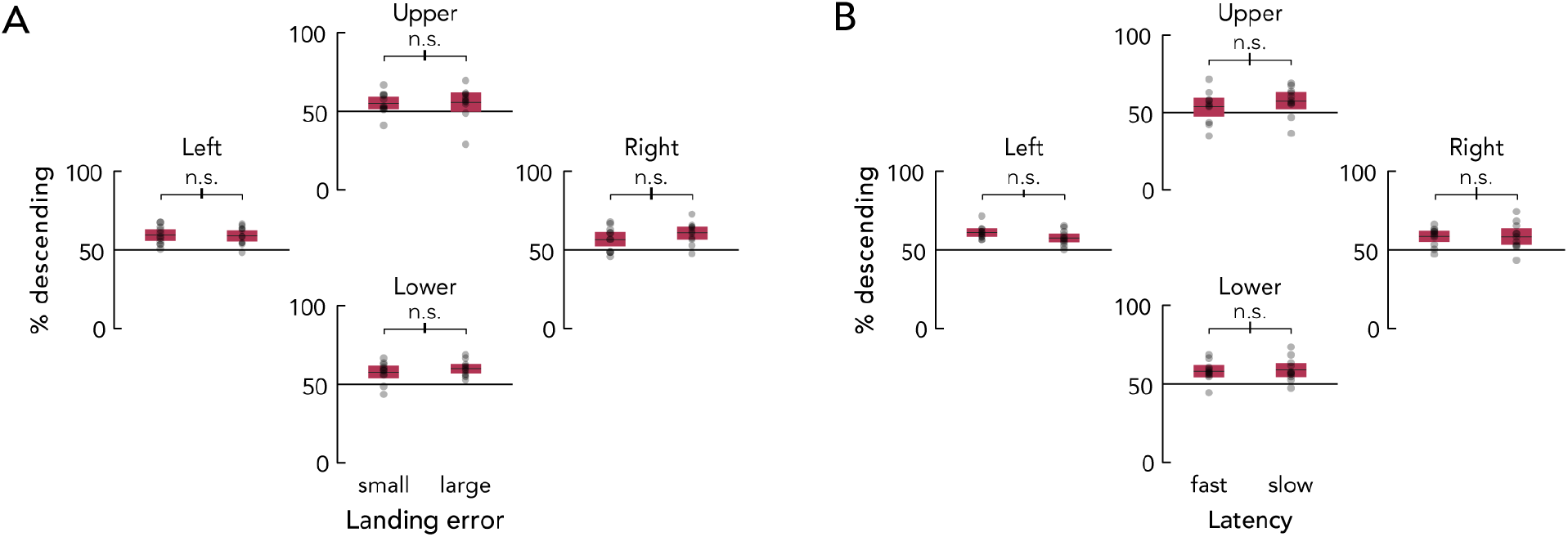
Influence of eye movement parameters on presaccadic perceptual enhancement. Percentage of trials contributing to staircase contrast decrement. Median-split for *valid* trials at each cardinal location with comparably small or large saccade landing error (**A**) and low or high saccade latency (**B**). Horizontal lines within each whisker plot indicate the group average. Bars depict the 95% confidence interval, black transparent dots represent individual observer data. Error bars on horizontal brackets show the SED.

## Discussion

This study reveals that visual performance asymmetries are not compensated by presaccadic attention. The benefit was neither more pronounced at the vertical than the horizontal meridian, nor was there any benefit at the upper vertical meridian, where contrast sensitivity is the worst. Thus, the intrinsic perceptual limitation at the upper vertical meridian observed during fixation (Carrasco et al., 2001; Cameron et al., 2002; Greenwood et al., 2017; Himmelberg et al., 2020; Barbot et al., 2021) is rigid, and cannot be mitigated even by the typically robust effect of presaccadic attention.

This impervious constraint might be explained by anatomical constraints in the retina and visual cortex. There are similar polar angle asymmetries in the density of photoreceptor cones and midget retinal ganglion cells (Curcio & Allen, 1990; Song et al., 2011; Watson, 2014), for which density is lowest for cells sampling along the upper vertical meridian of the visual field. However, a computational observer model has shown that these retinal asymmetries only account for a small proportion of behavioral contrast sensitivity asymmetries (Kupers et al., 2019, 2020). These asymmetries also exist, and are greatly amplified, in the distribution of cortical surface area in the primary visual cortex, where there is substantially less surface dedicated to processing the upper vertical meridian than the lower vertical meridian (Benson et al., 2021; Himmelberg et al., 2021a,b). Thus, there is a reduction in neural resources allocated to the upper vertical meridian that could likely explain poorer visual performance. Our results show that this lack of neural resources cannot be compensated for by pre-saccadic attention. The present findings demonstrate that the visual and neural circuitry underlying performance fields impose such pronounced limitations on visual processing that they may be very unlikely to be overcome.

Perceptual measurements are typically made at locations along the horizontal meridian. Likewise, studies investigating saccades (including those on presaccadic attention) typically rely on horizontal eye movements (e.g., Deubel & Schneider, 1996; Rolfs & Carrasco, 2012; Li et al., 2016, 2021b; Hanning et al 2019b; Ohl et al., 2017; Hübner & Schütz, 2021) or place the saccade targets around the visual field but do not investigate the effect of saccade direction (e.g., Kowler et al., 1995; Wollenberg et al. 2018; Hanning et al. 2019a). However, the existence of perceptual performance fields, as well as the current presaccadic data, question the generalizability of these findings and call for a systematic study of eye movements not only across eccentricity but also around the visual field.

Whereas in the present study we were particularly interested in separately assessing presaccadic benefits and costs (relative to fixation) to answer our research question –can presaccadic attention compensate for the characteristic polar angle performance asymmetries–, Montagnini and Castet (2007) investigated spatiotemporal dynamics of presaccadic attention by assessing the difference between performance at the saccade target and the (opposite) non-target location. The authors reported better orientation discrimination performance at the lower than the upper visual field, but concluded that this difference is not affected by the planning of upward or downward saccades, i.e., that there is no interaction between saccade target location and the presaccadic attention effect. Yet, by collapsing three tested locations across each hemifield, they may have neutralized a differential effect of presaccadic attention on the vertical meridian asymmetry as observed in the present study, given that performance (during fixation) gradually decreases from the horizontal to the vertical meridian, to the extent that it is similar at the four intercardinal locations (Abrams et al., 2012; Barbot et al., 2021). These differences within the hemifields highlight the importance of separately assessing presaccadic attention at each location (rather than collapsing across a hemifield), and comparing its effect to performance during fixation, before concluding that there is no interaction between saccade direction and presaccadic attention.

The present results have significant implications for human factors and ergonomic applications. For instance, the resilience of perceptual asymmetries around the visual field, even when actively exploring our environment with eye movements, should be known and taken into consideration when optimizing displays to accurately and efficiently convey relevant task information –for example, those used by pilots, air traffic controllers, drivers, and radiologists.

To summarize, saccade preparation enhanced contrast sensitivity, but surprisingly not at the upper vertical meridian, where contrast sensitivity is poorest and hence could benefit the most. Consequently, instead of diminishing contrast sensitivity asymmetries around the visual field, presaccadic attention actually exacerbates them and modifies the shape of the visual performance field.

### Limitations of the study

The generalizability of our initial, counterintuitive findings has yet to be assessed. The extent of performance fields is influenced by stimulus parameters; the polar asymmetries become more pronounced with eccentricity, spatial frequency and number of distracters (Carrasco et al 2001; Baldwin et al., 2012; Himmelberg et al. 2020; Barbot et al. 2021). Thus, it would be important to evaluate whether the exacerbated asymmetries that we report here with presaccadic attention still emerge while manipulating these stimulus parameters. Moreover, we know that the HVA and VMA in contrast sensitivity (Abrams et al., 2012) and visual acuity (Barbot et al., 2021) emerge gradually with angular distance at isoeccentric, perifoveal locations. It would be important to evaluate whether the effects of presaccadic attention not only differ at the cardinal meridians but also at intercardinal locations. Lastly, because presaccadic attention benefits many dimensions in addition to contrast sensitivity (review: Li et al., 2021a), its effect on orientation and motion sensitivity, spatial resolution, and their appearance should be assessed around the visual field.

## Materials and methods

### Observers

The sample size was determined based on previous work on presaccadic attention (Li et al., 2016; Ohl et al., 2017; Hanning et al., 2019a,b) and visual performance fields (Himmelberg et al., 2020; Barbot et al., 2021; Purokayastha et al., 2021). We report data of eleven observers (6 female, aged 19–32 years, two authors: NMH and MMH). All had normal or corrected-to-normal vision, provided written informed consent, and (except for two authors) were naive to the purpose of the experiment. Three additional observers were not considered in the final analysis because they did not meet our inclusion criteria – (1) typical performance asymmetries during fixation and (2) typical presaccadic attention effect^1^ (note that none of them showed a presaccadic benefit at the upper vertical meridian). The protocols for the study were approved by the University Committee on Activities involving Human Subjects at New York University and all experimental procedures were in agreement with the Declaration of Helsinki.

### Setup

Observers sat in a dimly illuminated room with their head stabilized by a chin and forehead rest and viewed the stimuli at 57 cm distance on a gamma-linearized 20-inch ViewSonic G220fb CRT screen (Brea, CA, USA) with a spatial resolution of 1,280 by 960 pixels and a vertical refresh rate of 100 Hz. Gaze position of the dominant eye was recorded using an EyeLink 1000 Desktop Mount eye tracker (SR Research, Osgoode, Ontario, Canada) at a sampling rate of 1 kHz. Manual responses were recorded via a standard keyboard. An Apple iMac Intel Core 2 Duo computer (Cupertino, CA, USA) running Matlab (MathWorks, Natick, MA, USA) with Psychophysics (Brainard, 1997; Pelli, 1997) and EyeLink toolboxes (Cornelissen, 2002), controlled stimulus presentation and response collection.

### Experimental design

The experiment comprised two eye movement conditions and a fixation condition. Eye movement conditions (*valid* and *invalid*) were randomly intermixed within blocks, whereas the fixation condition (*baseline*) was run in separate experimental blocks. At the beginning of each trial, observers fixated a central white dot (~52 cd/m^2^; diameter 0.45° of visual angle) on gray background (~26 cd/m^2^). Four placeholders indicated the isoeccentric locations of the upcoming stimuli (and potential saccade targets) 8.5° left, right, above, and below fixation. Each placeholder comprised four black dots (~0 cd/m^2^, diameter 0.1°), forming the corners of a squared diamond (diameter 4.2°). Once we detected stable fixation within a 2.25° radius virtual circle centered on this fixation, the beginning of the trial was indicated by a sound.

In eye movement blocks (*valid* and *invalid* trials), after 700 ms fixation period, a central direction cue (blue line, ~4 cd/m^2^, length 0.45°) pointed to one of the four cardinal placeholders (randomly selected), cueing the saccade target. Observers were instructed to look as fast and precisely as possible to the center of the indicated placeholder. 100 ms after cue onset (i.e., within the movement latency – gaze still rests at fixation), a Gabor grating (tilted ±15° relative to vertical; spatial frequency 5 cpd; 2.8° diameter Gaussian envelope diameter, σ = 0.43°) appeared for 50 ms either at the cued saccade target (*valid* trials; 50%) or at the location opposing the saccade target (*invalid* trials; 50%). Gabor contrast was titrated using an adaptive psychometric “staircase” procedure (see *Titration procedure*) and thus varied from trial to trial. Together with the Gabor, three radial frequency patterns with no orientation information (same spatial frequency, envelope, and contrast as the Gabor) were presented at the other placeholders to avoid biasing eye movements to a single sudden-onset stimulus. 300 ms after stimuli offset (once the eye movement had been performed), the dots of one placeholder increased in size (diameter 0.16°), functioning as a response cue to indicate the location that had contained the Gabor patch. Observers indicated their orientation judgement via button press (clockwise or counterclockwise, two-alternative forced choice) and were informed that the orientation report was non-speeded. They received auditory feedback for incorrect responses.

Stimulus parameters and timing for the fixation blocks (*baseline* condition) were identical to the eye movement blocks, with one difference: two (rather than one) blue direction cue lines appeared, pointing to opposing locations (left and right or upper and lower, randomly selected). Participants were instructed to keep eye fixation. As in the eye movement blocks, the Gabor appeared at one of two possible locations (indicated by the two direction cues) – thus location uncertainty as to where the test Gabor would appear was constant across experimental conditions.

Observers performed 3 sessions of 3 experimental blocks each (one fixation block followed by two eye movement blocks). Each block comprised 144 trials. We monitored gaze position online and controlled for correct eye fixation, i.e. gaze remaining within 2.25° from the central fixation target until (a) response cue onset (fixation blocks) or (b) direction cue onset (eye movement blocks). Observers maintained precise eye fixation during the pre-cue interval in fixation trials (0.80°±0.089° average distance from fixation target center ±1 SEM) as well as eye movement trials (0.72±0.077°). Trials in which gaze deviated from fixation were aborted and repeated at the end of each block. In eye movement blocks we also repeated trials with too short (<150 ms) or long (>350 ms) saccade latency, or incorrect eye movements (initial saccade landing beyond 2.25° from the indicated target). We collected a total of 1296 trials per observer – 432 fixation (*baseline*) trials and 864 eye movement trials (432 *valid*, 432 *invalid*).

### Titration procedure

We titrated contrast separately for each experimental condition (*valid, invalid, baseline*) and cardinal location (left, right, upper, lower) with best PEST (Pentland, 1980), an adaptive psychometric procedure, using custom code (https://github.com/michaeljigo/palamedes_wrapper) that ran subroutines implemented in the Palamedes toolbox (Prins & Kingdom, 2018). We concurrently ran 36 independent adaptive procedures (3 for each condition-location combination) targeting 80% orientation discrimination accuracy throughout the experiment. One psychometric procedure comprised 36 trials. We calibrated each procedure by presenting fixed levels of contrast that spanned the range of possible values (1% - 100%) for the first 9 trials. To derive the contrast thresholds, we took the median across the last 5 trials of each individual staircase. Then, before averaging across the 3 staircases per condition-location combination, we excluded outliers (3.03% of all procedures) for which the derived threshold deviated more than 0.5 log-contrast units from the other thresholds of the respective condition-location combination.

### Data analysis

We computed contrast sensitivity for each condition-location combination as the reciprocal of the average contrast threshold (CS = 1 / threshold). To evaluate the effect of saccade precision and latency on visual performance at the upper vertical meridian in the *valid* condition, we conducted two median splits and computed the percentage of trials causing the staircase procedure to decrease the contrast for saccades with smaller vs. larger landing error (***Figure 5A***) and saccades with faster vs. slower latencies (***Figure 5B***). Note that across the compared conditions, an average 58.25% of *valid* trials decreased the staircase contrast. Had saccade landing precision or latency affected the presaccadic attention benefit, trials with smaller / larger landing errors or faster / slower saccade latencies would have differentially affected staircase direction. This was not the case (see main text). For the conducted repeated-measures ANOVAs in which the sphericity assumption was not met, we report Greenhouse-Geisser corrected *p*-values; all *p*-values of post-hoc comparisons were Bonferonni corrected for multiple-comparisons.

## Additional information and files

### Author contributions

Conceptualization and methodology: NMH, MC; Software: NMH; Investigation: NMH, MMH; Formal analysis: NMH; Visualization: NMH, MMH; Writing—original draft: NMH; Writing—review & editing: NMH, MMH, MC; Funding acquisition: NMH, MC.

### Data availability

Raw eye tracking and behavioral data are available from the OSF database URL:https://osf.io/9a36u/.

**Supplemental Video S1**

Demonstration of stimuli and experimental design. Shown are one *valid*, one invalid, and one *baseline* trial. For demonstration purposes, all three trials test contrast sensitivity at the upper vertical meridian (contrast here fixed to 35%).

**Supplemental Table S2.** Correlation between visual field asymmetries (*HVA, VMA*) during fixation and presaccadic benefits (*valid / baseline*) and costs (*invalid / baseline*), for each tested cardinal location.

|  | Benefits |  |  |  | Costs |  |  |  |
| --- | --- | --- | --- | --- | --- | --- | --- | --- |
|  | Left | Right | Upper | Lower | Left | Right | Upper | Lower |
| **HVA** | R = -0.42 | R = -0.46 | R = -0.08 | R = -0.58 ^ | R = 0.03 | R = 0.20 | R = -0.05 | R = 0.09 |
| **VMA** | R = -0.16 | R = 0.10 | R = -0.20 | R = 0.46 | R = -0.01 | R = -0.10 | R = -0.08 | R = 0.04 |

Note. R = Pearson correlation coefficient. All (but one ^, $p > 0.05$ ) $p > 0.1$

## Acknowledgements

This research was supported by National Institutes of Health National Eye Institute grant R01 EY019693 to MC and a Feodor Lynen Research Fellowship from the Alexander von Humboldt Foundation to NMH. We thank Luke Huzsar, Michael Jigo, and other members of the laboratory of MC, as well as Jan Kurzawski and Hsin-Hung Li for useful comments and discussions. The authors declare no conflict of interest.

## Footnotes

1 One observer did not show the characteristic performance asymmetries during the fixation baseline condition (sensitivity horizontal meridian > lower vertical meridian > upper vertical meridian). Another observer did not show higher contrast sensitivity for *valid* trials than *invalid* trials. A third observer was excluded due to technical issues with eye movement recording.

